# Anticorrelation between default and dorsal attention networks varies across default subsystems and cognitive states

**DOI:** 10.1101/056424

**Authors:** Matthew L. Dixon, Jessica R. Andrews-Hanna, R. Nathan Spreng, Zachary C. Irving, Kalina Christoff

## Abstract

Anticorrelation between the default network (DN) and dorsal attention network (DAN) is thought to be an intrinsic aspect of functional brain organization reflecting competing functions. However, the stability of anticorrelations across distinct DN subsystems, different contexts, and time, remains unexplored. Here we examine DN-DAN functional connectivity across six different cognitive states. We show that:(i) the DAN is anticorrelated with the DN core subsystem, but not with the two DN subsystems involved in mentalizing and mnemonic functions, respectively; (ii) DN-DAN interactions vary significantly across cognitive states; (iii) DN-DAN connectivity fluctuates across time between periods of anticorrelation and periods of positive correlation; and (iv) coupling between the frontoparietal control network (FPCN) and DAN predicts variation in the strength of DN-DAN anticorrelation across time. These findings reveal substantial variability in DN-DAN interactions, suggesting that these networks are not strictly competitive, and that the FPCN may act to modulate their anticorrelation strength.

## Introduction

The last decade has witnessed extraordinary interest and progress in network neuroscience”the understanding of how interconnected brain regions operate in concert as large-scale networks, and how these networks relate to healthy and pathological cognitive functioning (Buckner, Krienen, & Yeo, 2013; Bullmore & Sporns, 2009; M. Fox & Raichle, 2007; Medaglia, Lynall, & Bassett, 2015; Petersen & Sporns, 2015). Resting state functional connectivity (rs-FC) has emerged as a powerful, non-invasive tool for delineating the functional network architecture of the human brain. Correlated fluctuations in BOLD signal measured during the “resting state” are thought to reveal intrinsic networks that persists across time (Damoiseaux et al., 2006) and context (Cole, Bassett, Power, Braver, & Petersen, 2014; Smith et al., 2009) due to their presumed source in stimulus-independent brain activity reflecting the underlying polysynaptic structural neuroanatomy (M. Fox & Raichle, 2007; Van Dijk et al., 2010).

One of the most influential findings to emerge from network neuroscience is that the default network (DN) and dorsal attention network (DAN) are anticorrelated during rest (M. Fox et al., 2005; see also Fransson, 2005; Golland, Golland, Bentin, & Malach, 2008). The DN has been linked to internally directed processes including self-reflection, autobiographical memory, future event simulation, and spontaneous thought (Andrews-Hanna, Smallwood, & Spreng,2014; Buckner, Andrews-Hanna, & Schacter, 2008; Christoff, Irving, Fox, Spreng, & Andrews-Hanna, in press; Ellamil et al., 2016; K. Fox, Spreng, Ellamil, Andrews-Hanna, & Christoff, 2015; Raichle et al., 2001), whereas the DAN has been linked to externally directed processes including attending to, and acting on behaviorally-relevant stimuli that are present in the immediate perceptual environment (Buschman & Kastner, 2015; Corbetta & Shulman, 2002; Golland et al., 2007; Miller & Buschman, 2013; Serences & Yantis, 2006; Szczepanski, Pinsk, Douglas, Kastner, & Saalmann, 2013). Based on these findings, Fox et al. (2005) proposed that anticorrelation between the DN and DAN is an adaptive mechanism that prevents interference between internal and external information processing. This idea has been used to explain the origin of attentional lapses and behavioral variability in healthy adults (Keller et al., 2015; Kelly, Uddin, Biswal, Castellanos, & Milham, 2008; Weissman, Roberts, Visscher, & Woldorff, 2006), cognitive immaturity in children (Chai, Ofen, Gabrieli, & Whitfield-Gabrieli, 2014), and abnormal functioning in conditions such as ADHD (Sonuga-Barke & Castellanos, 2007). Although global signal regression can induce spurious anticorrelations when included as part of data preprocessing (Murphy, Birn, Handwerker, Jones, & Bandettini, 2009; Saad et al., 2012), DN-DAN anticorrelation is observed even without this step, suggesting that it is a true biological phenomenon (Chai, Castanon, Ongur, & Whitfield-Gabrieli, 2012; M. Fox, Zhang, Snyder, & Raichle, 2009).

While the idea that the DN and DAN are intrinsically anticorrelated has intuitive appeal, theoretical and empirical work suggests the possibility of a more variable relationship between these networks and the processes they support (E. Allen et al., 2014; Chang & Glover, 2010; Dixon, Fox, & Christoff, 2014; Kelly, et al., 2008; Meyer, Spunt, Berkman, Taylor, & Lieberman, 2012; Spreng et al., 2014; Summerfield, Lepsien, Gitelman, Mesulam, & Nobre, 2006). However, the potential variability of DN-DAN interactions has not been the primary focus of prior work, and to our knowledge, no studies have systematically investigated whether DN-DAN interactions differ across regions, cognitive states, and time. If observed, such variability would offer a novel perspective on large-scale network interactions underlying the capacity to process information about the internal and external environments.

Since the discovery of DN-DAN anticorrelation, developments in understanding the DN have now revealed that it is not a unitary entity, but rather, composed of three distinct subsystems (for a review see Andrews-Hanna, et al., 2014). Our first goal was to examine the extent to which anticorrelations are present for all three subsystems of the DN. Although it is too early to definitively characterize the function of each subsystem, preliminary evidence suggests:(1) a core subsystem involved in self-referential processing, including the construction of a temporally-extended self with attributes, preferences, and autobiographical details; (2) a dorsomedial subsystem involved in semantic processing and mentalizing (i.e., generating inferences about mental states including beliefs, desires, and intentions); and (3) a medial temporal subsystem involved in retrieving and binding together contextual details during the recollection of episodic memories and simulation of future events. Interestingly, studies have found coactivation of the DAN and dorsomedial subsystem during a social working memory task (Meyer, et al., 2012), and coactivation of the DAN and medial temporal subsystem during a memory-guided attention task (Summerfield, et al., 2006), raising the possibility that these subsystems may not be antagonistic (anticorrelated) with the DAN. Indeed, learning often requires a synergy between perceptual and memory processes (Chun & Turk-Browne, 2007; Hasselmo & McGaughy, 2004), and mental state inferences often draw upon perceptual input (e.g., facial expressions) (Baron-Cohen, Wheelwright, Hill, Raste, & Plumb, 2001). Discerning the nature of functional interactions between the DAN and the distinct DN subsystems would provide critical information about the cognitive processes that may or may not be inherently competitive.

A second goal of the present study was to systematically examine the stability of anticorrelations across time and different cognitive states. Evidence is accumulating that the strength and topography of functional connectivity patterns dynamically change across time and cognitive state (E. Allen, et al., 2014; Braun et al., 2015; Cole et al., 2013; Davison et al., 2015; Geerligs, Rubinov, Cam, & Henson, 2015; Gonzalez-Castillo et al., 2015; Hermundstad et al., 2014; Hutchison, Womelsdorf, Gati, Everling, & Menon, 2013; Krienen, Yeo, & Buckner, 2014; Mennes, Kelly, Colcombe, Castellanos, & Milham, 2013; Shirer, Ryali, Rykhlevskaia, Menon, & Greicius, 2012). It is possible that anticorrelations are specifically related to the cognitive state elicited by rest, that is, spontaneous thoughts of current concerns, past events, and future plans (Andrews-Hanna, 2012; Delamillieure et al., 2010). A recent study observed that the default network can be engaged during an externally oriented working memory task when participants leveraged prior knowledge of the stimuli to complete the task (Spreng, et al., 2014). This finding suggests there may be task conditions that afford greater cooperation between the DN and DAN. Furthermore, there is some evidence that anticorrelations may vary across time even during rest (E. Allen, et al., 2014; Chang & Glover, 2010). Here, we investigated possible contextual and temporal variability of DN-DAN interactions by examining their relationship across time and different cognitive states within the same participants.

Our third goal was to examine whether the frontoparietal control network (FPCN) plays a role in modulating DAN-DN functional connectivity. Recent theoretical and empirical work suggests that the FPCN contributes to the control of internal and external attention by modulating processing in specific regions based on current goals (Dixon, et al., 2014; Gao & Lin, 2012; Smallwood, Brown, Baird, & Schooler, 2012; Spreng, Stevens, Chamberlain, Gilmore, & Schacter, 2010; Vincent, Kahn, Snyder, Raichle, & Buckner, 2008). This idea is consistent with the role of the FPCN in executive control (Duncan, 2010; Miller & Cohen, 2001), the extensive functional interconnections linking the FPCN to the DN and DAN (Spreng, Sepulcre, Turner, Stevens, & Schacter, 2013), and evidence that the FPCN flexibly couples with other networks including the DN and DAN and drives large-scale network reconfiguration based on task demands (Braun, et al., 2015; Cole, et al., 2013; Fornito, Harrison, Zalesky, & Simons, 2012; Gao & Lin, 2012; Spreng, et al., 2010). In healthy older adults, reduced anticorrelation between the DN and DAN has been observed (Keller, et al., 2015; Spreng, Stevens, Viviano, & Schacter, in press), and may relate to a shift towards greater between-network connectivity with advancing age, particularly with the FPCN (Grady, Sarraf, Saverino, & Campbell, 2016). However, it is currently unknown whether there is a direct relationship between patterns of FPCN connectivity and the strength of DN-DAN anticorrelation. Here we aimed to assess this possibility.

To examine these three questions, we used fMRI in conjunction with functional connectivity and machine learning classification analyses. We monitored brain activation dynamics during six conditions (see **Experimental Procedures**):(i) rest; (ii) movie viewing; (iii) analysis of artwork; (iv) social preference shopping task; (v) evaluation-based introspection; and (vi) acceptance-based introspection. Because these conditions differ from traditional cognitive tasks, we refer to them as cognitive states or contexts, rather than tasks. They were designed to elicit mental states that resemble those frequently experienced in everyday life. Furthermore, they were designed to include a combination of external and internal processing requirements. Each condition elicited a continuous mental state and did not require any responses, making each similar to rest. All data underwent the same preprocessing procedure typically used with resting state fMRI that does not rely upon global signal regression (Whitfield-Gabrieli & Nieto-Castanon, 2012).

## Results

**Presence of Anticorrelations Across DN Subsystems**. We first examined whether the DAN is anticorrelated with all three DN subsystems during rest. To explore these interactions in relation to well-established network boundaries, we used regions of interest (ROIs) created by Yeo and colleagues (Krienen, et al., 2014; Yeo, Tandi, & Chee, 2015) based on their 17-network parcellation derived from the data of 1000 participants (Yeo et al., 2011) (see **Supplemental Experimental Procedures**; Figure 1A; **Figure S1**).We extracted the mean activation timeseries from each of 32 ROIs spanning the DAN and three DN subsystems, and calculated the timeseries correlation between pairs of regions belonging to the DN and DAN. We then computed the average strength of connectivity between the DAN and each DN subsystem. The results demonstrated that connectivity strength significantly varied across subsystems [*F*(2, 46) = 17.78, *p* <.001]. The DAN was anticorrelated with the Core subsystem (r = − .13), but showed little to no anticorrelation with the dorsomedial (*r* = − .01) or medial temporal subsystems (*r* = − .04) (Figure 1B). Consistent with this, whole-brain voxel-wise analyses revealed that DAN seed regions exhibited negative connectivity with voxels primarily located within the borders of the Core subsystem (Figure 1C). To further examine these relationships, we created connectivity fingerprints for DAN ROIs and observed that negative connectivity was mainly observed with Core subsystem regions (Figure 1D). These findings reveal that anticorrelations are spatially specific. For example, anticorrelation is robust for the rostromedial prefrontal cortex but weak for the adjacent dorsomedial prefrontal cortex, and robust for the posterior inferior parietal lobule, but weak for the adjacent temporoparietal junction. Moreover, region aMT of the DAN did not exhibit anticorrelation with any DN regions. Together, these findings demonstrate regional variability in DN-DAN interactions, with little evidence of an inherent competition between the dorsomedial and medial temporal subsystems and the DAN.

**Figure 1.**
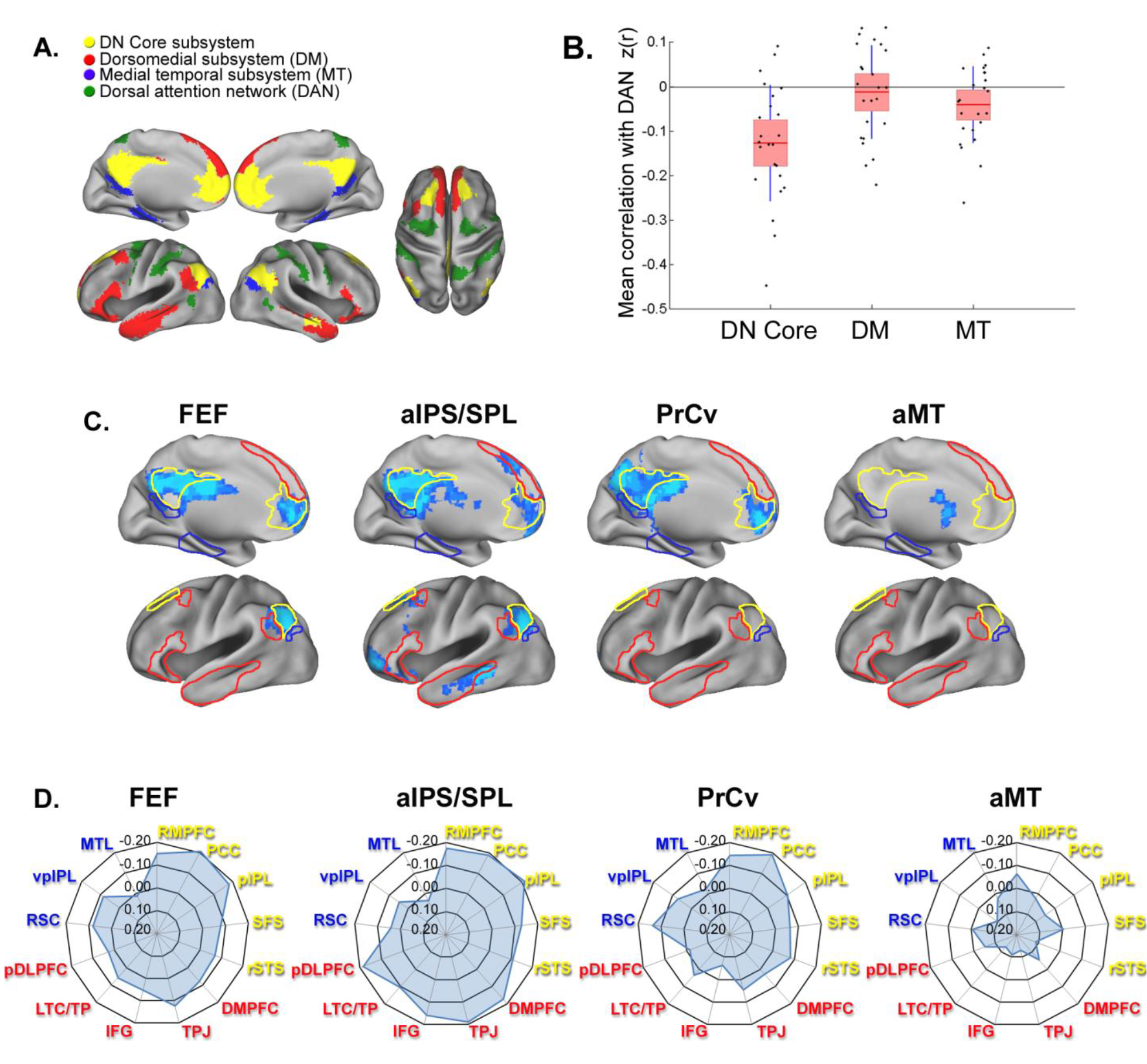
Anticorrelations as a function of DN subsystem. (A) Networks from Yeo et al. (2011) used for ROIs. (B) Mean correlation between the DAN and each DN subsystem. Data for each participant (black dots), with mean (red line), 95% CI (red shaded area) and 1 SD (purple lines). Anticorrelation was stronger for the Core subsystem relative to the dorsomedial and medial temporal subsystems [*t*(23) = 5.59, *p* <.001 and *t*(23) = 4.02, *p* = .001, respectively]. (C) Seed-based connectivity analyses showing negative connectivity with DAN regions (*Z* >2.57, *p* <.05 FDR corrected for cluster extent), with the borders of each DN subsystem highlighted. DAN seeds:FEF, frontal eye fields; aIPS/SPL, anterior intraparietal sulcus/superior parietal lobule; PrCv, ventral precentral cortex; aMT, anterior middle temporal region. Left hemisphere data is presented (see **Figure S2** for right hemisphere data). (D) Functional connectivity fingerprints for each DAN region. Core subsystem:RMPFC, rostromedial prefrontal cortex; PCC, posterior cingulate cortex; pIPL, posterior inferior parietal lobule; SFS, superior frontal sulcus; rSTS, rostral superior temporal sulcus. DM subsystem:DMPFC, dorsomedial prefrontal cortex, TPJ, temporoparietal junction, TP/LTC, temporopolar cortex/lateral temporal cortex; IFG, inferior frontal gyrus, pDLPFC, posterior dorsolateral prefrontal cortex. MT subsystem:MTL, medial temporal lobe; RSC, retrosplenial cortex; vpIPL, ventral posterior inferior parietal lobule.

**Stability of Anticorrelations Across Cognitive States**. Next, we examined whether anticorrelations exhibit stability across different cognitive states. Prior work has examined the stability of connectivity patterns by computing the correlation between context-specific connectivity matrices (Cole, et al., 2014; Geerligs, et al., 2015; Krienen, et al., 2014). Strong correlations imply that connectivity patterns are highly similar across contexts, thus suggesting stability. Here, we adopted this approach, but focused specifically on DN-DAN connections rather than whole-brain connectivity patterns (Figure 2A). As illustrated in Figure 2B, the similarity between DN-DAN connectivity patterns across different cognitive contexts was modest. Critically, across-context similarity was significantly lower than within-context similarity” that is, the similarity of DN-DAN connectivity from the first half to the second half of each context. This was the case when considering all DAN-DN pairwise connections [paired *t*-test:*t*(23) = 10.46, *p* <.001], and when breaking down the analysis by DN subsystem [Core:*t*(23) = 7.84,*p* <.001; dorsomedial:*t*(23) = 5.61,*p* <.001; medial temporal:*t*(23) = 9.35,*p* <.001]. The sizable difference between within- and across-context similarity reveals a substantial effect of context on DN-DAN connectivity. Importantly, this was not due to the separation of contexts in time; nearly identical results were obtained when comparing connectivity during one context to connectivity during the immediately preceding context (**Figure S3**). These findings reveal that anticorrelation strength varies considerably across different cognitive states.

**Figure 2.**
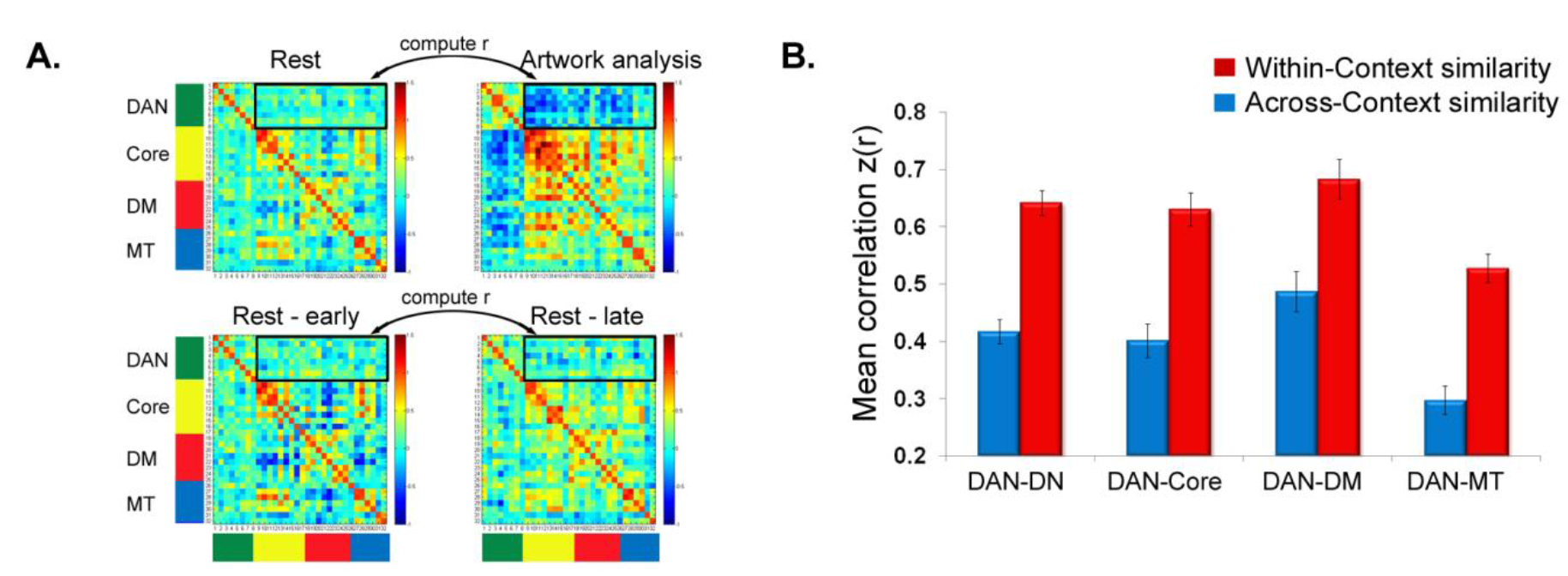
Comparison of within- and across-context similarity of DN-DAN connectivity. (A) Example of the analysis approach for one participant. We extracted DN-DAN correlation values (highlighted by the black box), and then calculated the correlation between the vector of connectivity values for each pair of cognitive contexts, and between the vector of connectivity values for the early and late period within each context. (B) Mean within- and across-context similarity of anticorrelations. DAN, dorsal attention network; DN, entire default network; DM, dorsomedial subsystem; MT, medial temporal subsystem.Error bars reflect within-subject SEM (Loftus & Masson, 1994).

To further interrogate whether DN-DAN connectivity patterns flexibly reconfigure in each cognitive context, we examined whether a support vector machine (SVM) classifier could accurately distinguish each pair of cognitive states solely on the basis of DN-DAN connectivity patterns. Above chance-level accuracy would imply distinct connectivity patterns in each context. The SVM was fed training data (a vector consisting of all DN-DAN correlations) and learned a model that maximized the separation of two cognitive states (e.g., rest and movie viewing) in multidimensional space, based on the pattern of features defining each context. The SVM then used its model of the training data to predict the labels of new data. Classifier accuracy was determined using leave-one-out cross validation, and statistical significance was established using permutation testing. As depicted in Figure 3, the SVM achieved classification accuracy that was considerably above chance-level (*ps* <.05) in 12/15 comparisons. This suggests that the SVM classifier could distinguish each pair of cognitive states solely on the basis of DN-DAN connectivity patterns, thereby implying a relatively unique configuration of DN-DAN interactions within each context that was reliable across subjects.

**Figure 3.**
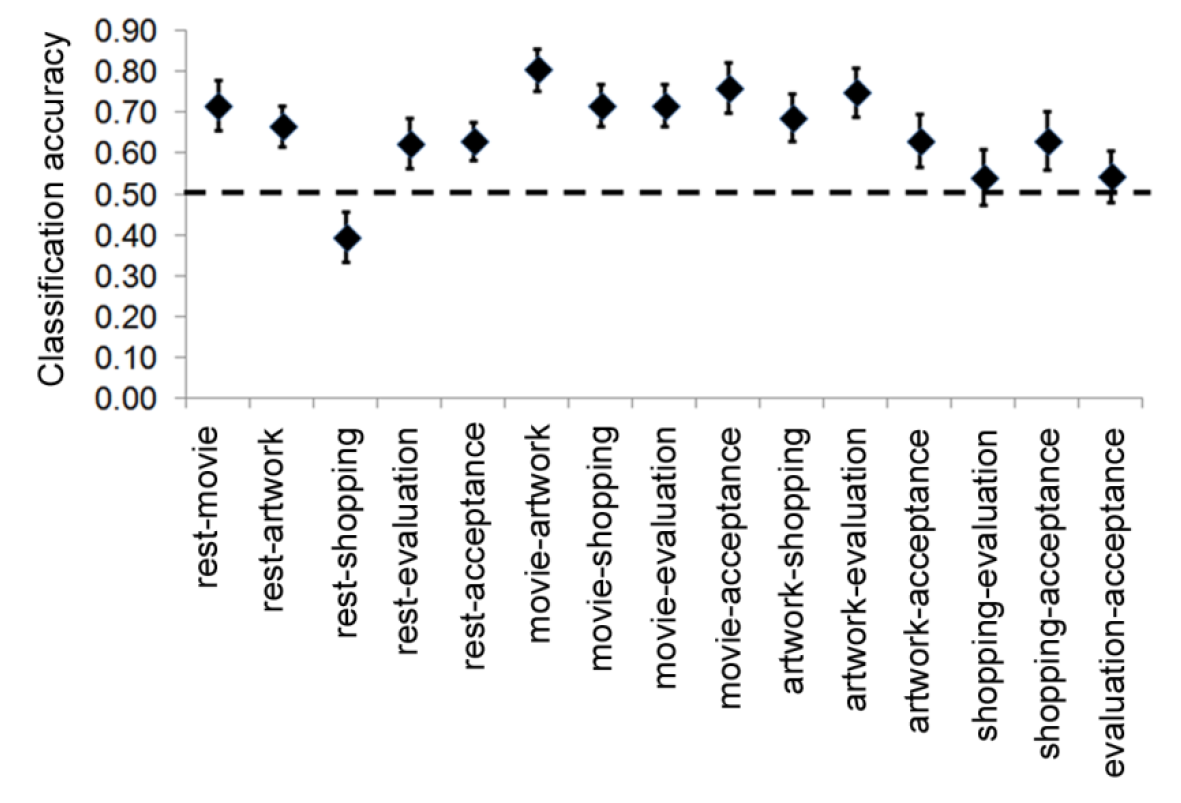
Accuracy of the pattern classifier in distinguishing each pair of cognitive contexts. Classification accuracy was significantly above chance level (*p* <.05) in all cases except for the Rest-Shopping, Shopping-Evaluation, and Evaluation-Acceptance comparisons. Error bars reflect between-subject SEM.

To provide more detail regarding the direction of changes in DN-DAN connectivity across different cognitive states, we conducted whole-brain seed-based analyses. The results demonstrated that anticorrelations flexibly increased or decreased in different cognitive contexts relative to rest (Figure 4; *Z* >2.57, *p* <.05 FDR corrected for cluster extent). A pair of DN-DAN regions could exhibit anticorrelation in one context, but no anticorrelation or even positive connectivity in other contexts. Moreover, it is apparent that the effect of context is region-specific:some DN-DAN connections exhibit little contextual variance, whereas other connections shift considerably. This suggests that a broad interpretation of cognitive functions based on a single summary measure of DN-DAN interactions will not accurately capture the true nature of their relationship. Importantly, control analyses ruled out the possibility that the effect of context was driven by motion (see **Supplemental Experimental Procedures**).

**Figure 4.**
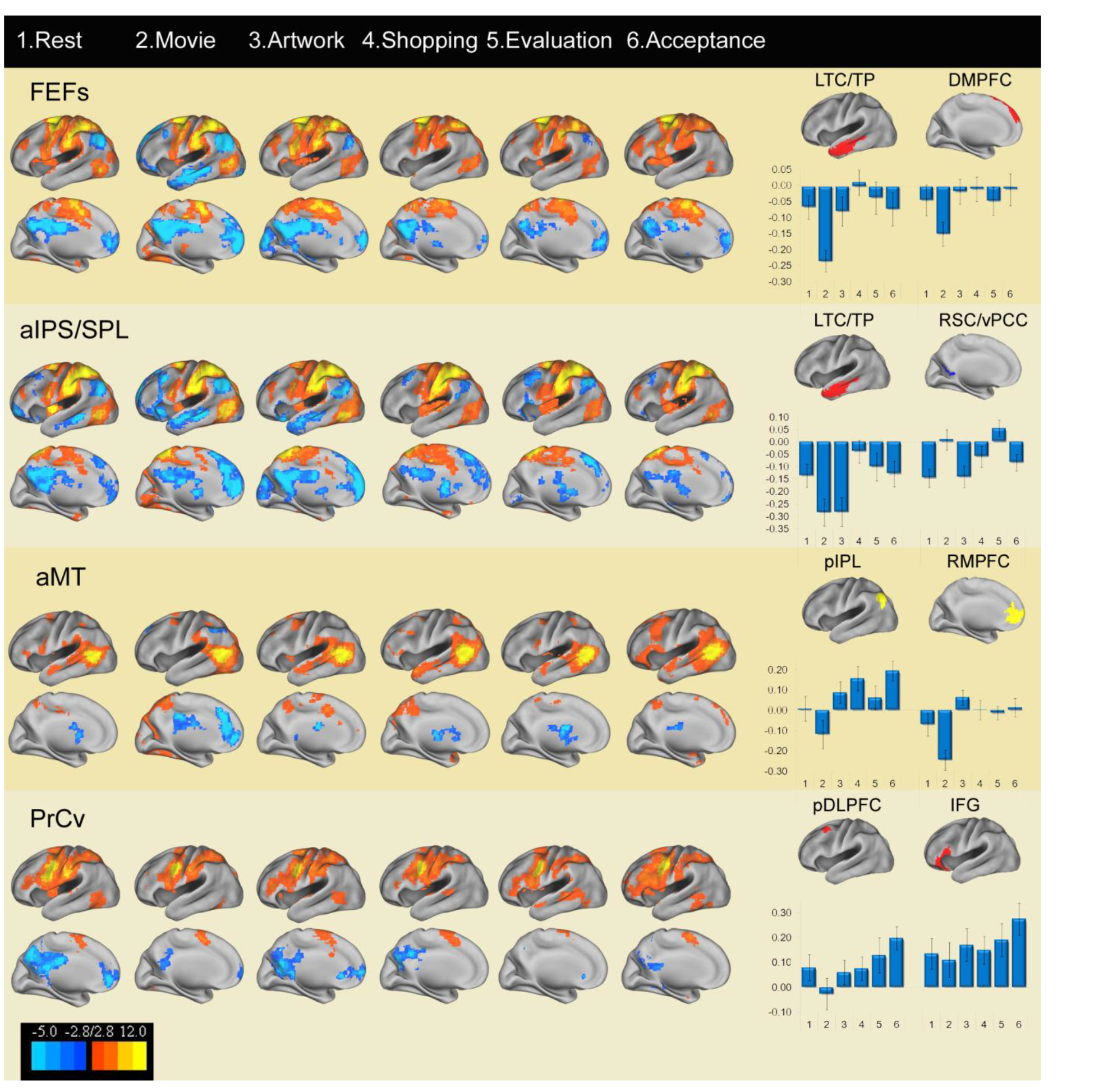
Whole-brain seed-based analyses. Positive and negative functional connectivity for each DAN seed region and context. Anticorrelations flexibly increase and decrease in different cognitive contexts relative to rest (*Z* >2.57, *p* <.05 FDR cluster corrected). Right panel:mean connectivity strength, *z*(*r*), for specific pairs of DN-DAN ROIs for each context. Results for left hemisphere presented (see Figure S4 for right hemisphere data). Colour bar shows *t*-values.

**Stability of Anticorrelations Across Time**. DN-DAN interactions are generally summarized as a single correlation value reflecting connection strength across a long period of time (e.g., 5-10 minutes). While useful, this approach cannot reveal potential temporal variation in DN-DAN interactions. The contextual variance reported above raises the possibility that anticorrelation strength is influenced by an individual's current mental state, and could potentially shift across time even during rest in accordance with changing mental content. We investigated time-resolved DN-DAN connectivity using a 60-second sliding window approach (Hutchison et al., 2013). Prior work has shown that functionally-relevant connectivity patterns can be isolated from ∼ 60 seconds of data (Gonzalez-Castillo, et al., 2015; Leonardi & Van De Ville, 2015; Liegeois et al., 2015; Shirer, et al., 2012). For each participant, we computed average DN-DAN connectivity within each window during rest, and then calculated the percentage of windows during which anticorrelation was present. The results demonstrated considerable temporal variability, with the DN and DAN alternating between anticorrelated and uncorrelated/positively correlated states (Figure 5A). On average, the DAN was anticorrelated in 67.09%of windows with the Core subsystem, in 52.75% of windows with the dorsomedial subsystem, and in 56.16% of windows with medial temporal subsystem (Figure 5B). Anticorrelation was observed in a greater number of windows for the Core subsystem relative to the dorsomedial and medial temporal subsystems (paired *t*-test:*t*(23) = 4.38,*p* <.001 and *t*(23) = 3.68,*p* = .001, respectively), recapitulating the distinction between the subsystems observed in the standard analysis. However, even in the case of the Core subsystem there were frequent shifts away from an anticorrelated state.

**Figure 5.**
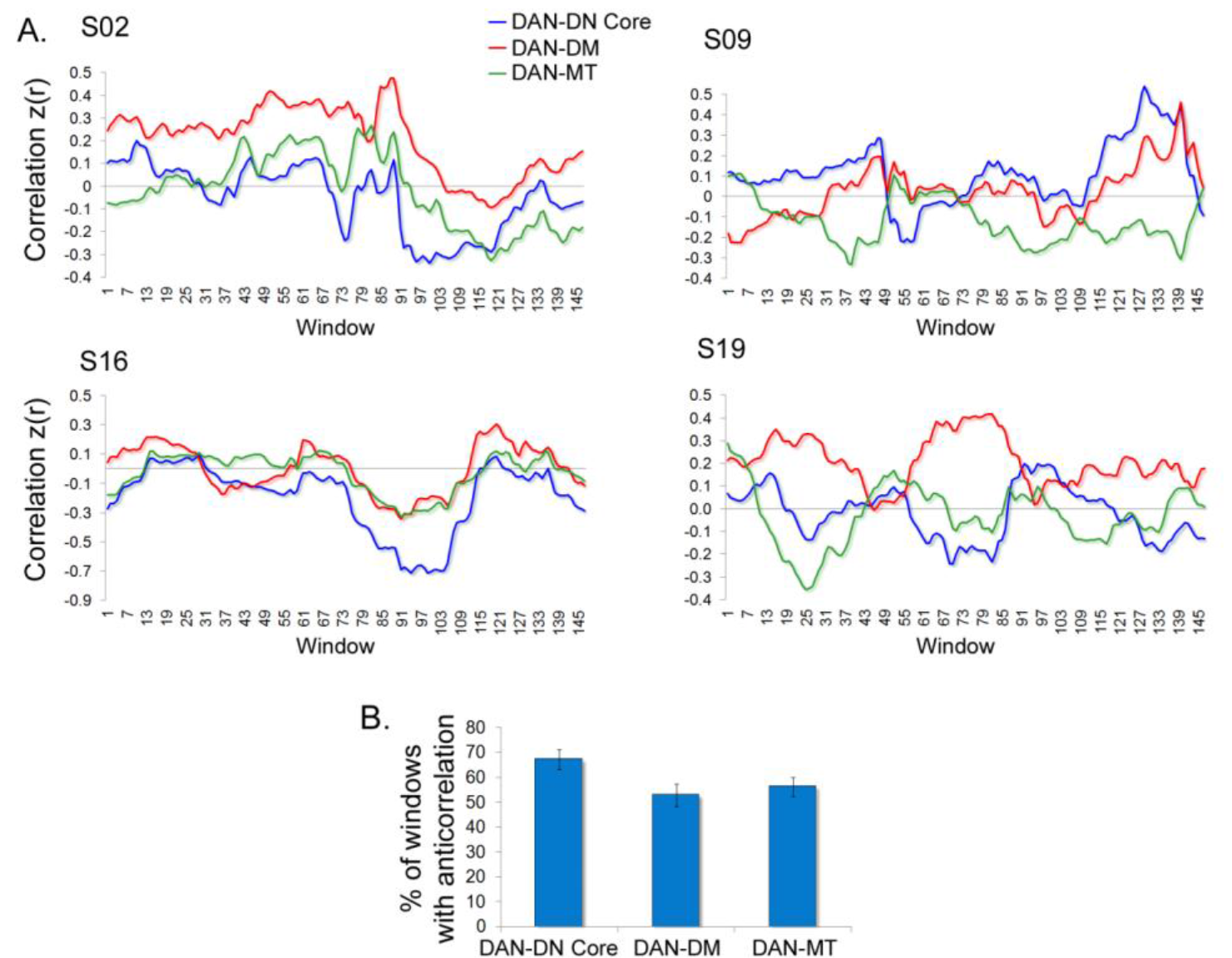
Temporal variability in DN-DAN interactions during rest. (A) Data for four randomly chosen example participants demonstrating average correlation strength between the DAN and each DN subsystem within successive 60-second windows. For each subsystem, connectivity with the DAN alternated between periods of anticorrelation and periods of positive correlation. (B) Percentage of windows during which DN-DAN connectivity was anticorrelated, averaged across all participants. DM, dorsomedial subsystem; MT, medial temporal subsystem. Error bars represent between-subject SEM.

**Frontoparietal Control Network Connectivity and Anticorrelations**. Having demonstrated that anticorrelations vary in strength across time, we next sought to identify potential modulatory influences that may help to explain this variance. Based on evidence that the frontoparietal control network (FPCN) (Figure S5) flexibly couples with the DN and DAN, we hypothesized that it may play a role in modulating the strength of anticorrelations. Because only the Core subsystem showed evidence of reliable anticorrelation with the DAN, we focused on this subsystem in the following analysis. We adapted recently derived methods from dynamic network science (Bassett, Wymbs, Porter, Mucha, & Grafton, 2014; Davison, et al., 2015) to provide a hypothesis-driven examination of whether changes across time in the strength of DAN-Core anticorrelations were correlated with temporal fluctuations in FPCN-DAN and FPCN-Core connectivity patterns. Within each 60-second window, we computed the average strength of FPCN-DAN, FPCN-Core, and DAN-Core connectivity, providing a timeseries of between-network connectivity values. We then computed the correlation between the timeseries to examine the co-evolution of temporal variation in FPCN connectivity patterns and anticorrelation strength. The results demonstrated a strong positive correlation between temporal fluctuations in FPCN-DAN connectivity and DAN-Core connectivity”a relationship that held in every context (Figure 6A and 6B). Periods of time characterized by stronger negative FPCN-DAN coupling were associated with stronger DAN-Core anticorrelation, whereas periods of little to no FPCN-DAN coupling were associated with weaker DAN-Core anticorrelation. Importantly, this result cannot be attributed to a general effect such as global fluctuations in BOLD signal, as temporal fluctuations in FPCN-Core connectivity demonstrated the opposite pattern, at least during rest and movie viewing (Figure 6A and 6B). Periods of time characterized by stronger positive FPCN-Core coupling were associated with stronger DAN-Core anticorrelation during these conditions. Finally, in four of six contexts, we found that periods of stronger positive FPCN-Core coupling were associated with stronger FPCN-DAN anticorrelation (Figure S6), supporting the idea of a triadic relationship. Thus, when FPCN signal was more coupled with the Core it was more anticorrelated with the DAN, and this was accompanied by stronger DAN-Core anticorrelation (Figure 6C). These systematic relationships are consistent with the idea that the FPCN may potentially modulate the strength of anticorrelations via interactions with the DN and DAN.

**Figure 6.**
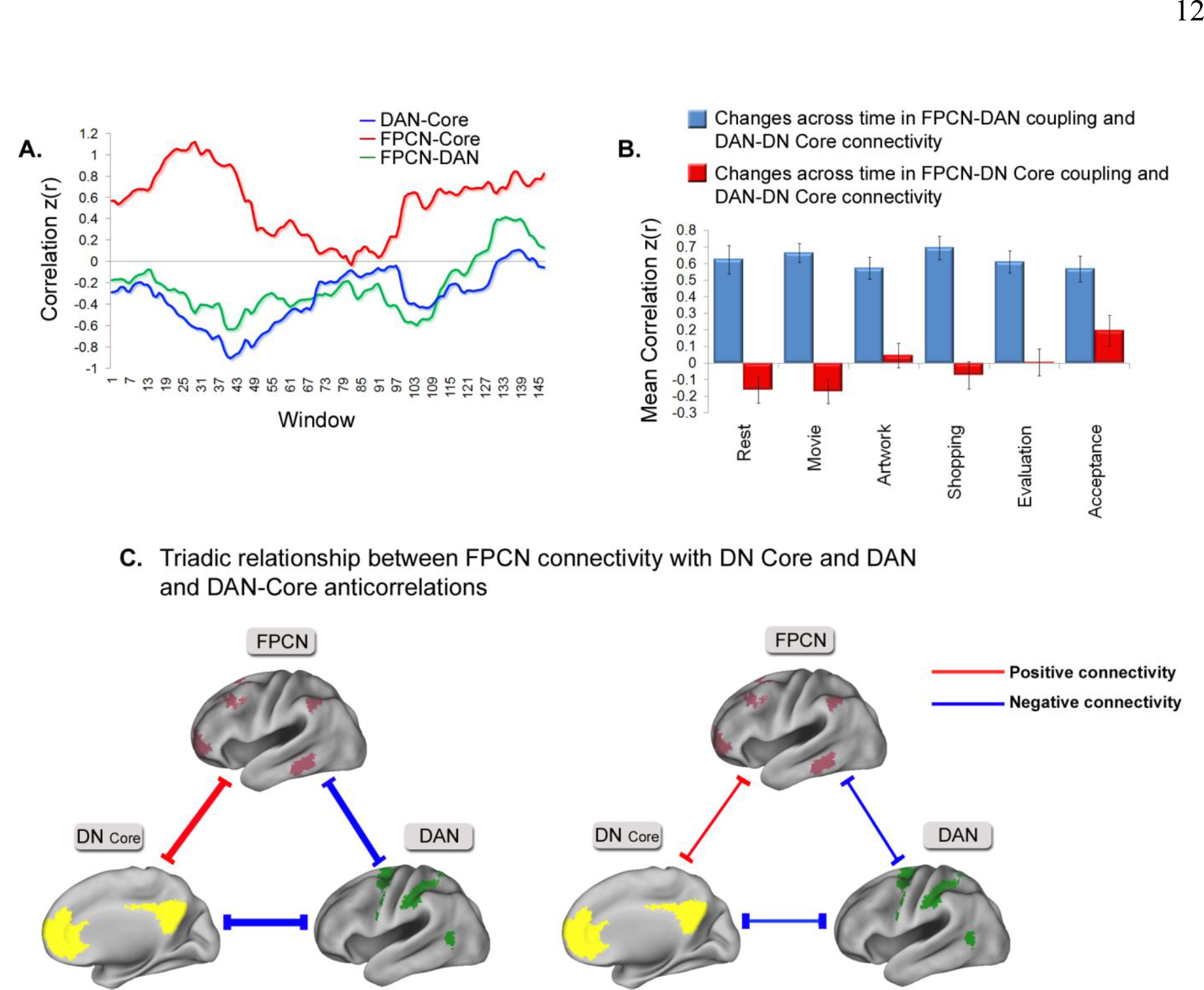
Relationship between FPCN connectivity and DN-DAN anticorrelation. (A) Data for an example participant showing that fluctuations across time in FPCN connectivity are associated with the strength of DAN-Core anticorrelation. (B). Temporal fluctuations in FPCN-DAN coupling are associated with DAN-Core anticorrelation strength in every context (all *p* <.001). Temporal fluctuations in FPCN-Core coupling are associated with DAN-Core anticorrelation strength during rest (*p* = .055), movie viewing (*p* = .028), and acceptance-based introspection (*p* = .026), but not in other contexts (*ps* >.05). Error bars reflect between-subject SEM. (C) Schematic illustration of the relationship between FPCN connectivity and DN-DAN anticorrelation. Width of the lines represents connectivity strength.

## Discussion

Delineating the nature of functional interactions between the DN and DAN is critical for understanding how attention is efficiently allocated to internal thoughts and external perceptual information. Although prior work suggests that the DN and DAN are intrinsically anti correlated, our findings, using well-established network boundaries (Yeo, et al., 2011), provide compelling evidence that DN-DAN interactions are more variable than previously thought:(i) the DAN was not anticorrelated with all three DN subsystems; (ii) DN-DAN interactions flexibly reconfigured across different cognitive states; and (iii) DN-DAN connectivity fluctuated across time between periods of anticorrelation and periods of positive correlation. Notably, we did observe one consistent relationship:temporal fluctuations in FPCN-DAN coupling were correlated with changes across time in the strength of DN-DAN anticorrelation within every context, suggesting that the FPCN may play a role in modulating the strength of anticorrelations. Together, these findings suggest that the DN and DAN and the functions they support are not strictly antagonistic.

**Anticorrelations Vary Across DN Subsystems**. The DAN exhibited modest anticorrelation with the Core subsystem, but little to no anticorrelation with the dorsomedial and medial temporal subsystems. Notably, Fox et al.'s (2005) original report of DN-DAN anticorrelation was based on seed regions located within the Core subsystem. We therefore replicate Fox et al.'s (2005) original results, while providing evidence against the hypothesis that the DN as a whole is competitive with the DAN. The relationship between these networks has often been interpreted in terms of competing internally- and externally-directed functions, however, our results highlight the complex nature of interactions between the DN and DAN, and do not support the idea of a strict antagonism between internal and external processing (Dixon, et al., 2014). Rather, antagonism is specifically related to the function of the DN Core subsystem. The DN Core is invariably recruited when individuals reflect on the self as an object of awareness with particular goals, attributes, and a linear narrative that connects past, present, and future experience (Denny, Kober, Wager, & Ochsner, 2012; Farb et al., 2007; Murray, Schaer, & Debbane, 2012; Schmitz & Johnson, 2007; Wagner, Haxby, & Heatherton, 2012), consistent with the idea that it contributes to an autobiographical mode of self processing (Araujo, Kaplan, Damasio, & Damasio, 2015; Christoff, Cosmelli, Legrand, & Thompson, 2011; Farb, et al., 2007; Gallagher, 2000). One possibility is that anticorrelations shield this type of abstract self-reflection from potential interference by perceptual processes supported by the DAN.

In agreement with our results, numerous lines of evidence suggest that mentalizing and mnemonic processes associated with the dorsomedial and medial temporal subsystems are not inherently antagonistic with perceptual processes associated with the DAN (Dixon, et al., 2014). For example, memory can facilitate the deployment of attention to the external environment (e.g., remembering where one last put the car keys) and this is subserved by co-activation of medial temporal and DAN regions (Summerfield, et al., 2006). Similarly, another study found that working memory performance was facilitated for famous relative to unfamiliar faces, and this was accompanied by medial temporal subsystem activation, consistent with the idea that mnemonic representations can facilitate perceptual encoding when it is congruent with task demands (Spreng, et al., 2014). Furthermore, during the encoding of new information, medial temporal regions decouple from other DN regions (Huijbers, Pennartz, Cabeza, & Daselaar,2011), and become more sensitive to afferent sensory input, as a result of acetylcholine's modulatory influence on medial temporal lobe circuit dynamics (Hasselmo & McGaughy, 2004). Finally, in the case of rest, the spontaneous reactivation of information stored in memory may in some cases lead to an autobiographical stream of thought that becomes elaborated upon by the Core subsystem, but in other cases may trigger a sensorimotor stream of thought (e.g., an imagined interaction with the environment) that may elicit cooperative medial temporal subsystem-DAN dynamics. Accordingly, the medial temporal subsystem may go in and out of phase with the DAN depending on whether mnemonic and perceptual processes pertain to the same or different goals, thus resulting in uncorrelated activation on average.

Similarly, mentalizing and perceptual processing may sometimes operate in concert, as perception of body language, facial expression, and eye-gaze often inform the inferences we make about others’ thoughts, and vice versa (Baron-Cohen, et al., 2001). Supporting this idea, coactivation of the DAN and dorsomedial subsystem is observed when individuals view dynamic animations and attend to the social intentional meaning of the movements (Tavares, Lawrence, & Barnard, 2008). Thus, mentalizing and memory processes are sometimes, but not always associated with perceptual decoupling (Schooler et al., 2011; Smallwood, et al., 2012). The brain has limited attentional resources, and consequently, has difficulty performing more than one goal at a time (Marois & Ivanoff, 2005). When mentalizing and mnemonic processes can be linked to perceptual processing in service of a unified goal, there may be little to no interference, but when they pertain to different goals (e.g., during mind wandering) they are likely to compete (Dixon, et al., 2014). In line with this, our dynamic connectivity analysis revealed that the DAN flexibly shifts between periods of anticorrelation and periods of positive correlation with the dorsomedial and medial temporal subsystems even during rest. This may reflect the exploration of frequently occurring network states.

**Contextual and Temporal Variability of Anticorrelations**. A burgeoning literature has revealed context-dependent FC patterns, with an emerging picture of the brain as a dynamic evolving system that flexibly adapts to changes in internal and external states (E. Allen, et al., 2014; Braun, et al., 2015; Cole, et al., 2013; Davison, et al., 2015; Geerligs, et al., 2015; Gonzalez-Castillo, et al., 2015; Krienen, et al., 2014; Mennes, et al., 2013; Milazzo et al., 2014; Shirer, et al., 2012; Spreng, et al., 2010). Connectivity patterns have been linked to individuals' mental states (Andrews-Hanna, Reidler, Huang, & Buckner, 2010; Doucet et al., 2012; Gorgolewski et al., 2014), and flexibility appears to be adaptive, given that it correlates with task performance (Braun, et al., 2015; Hermundstad, et al., 2014). Building upon this work, we report convergent findings revealing that anticorrelations exhibit variability across different cognitive states.

Our similarity analysis revealed little stability in DAN-DN connectivity across different cognitive contexts. Consistent with this, a prior study found that anticorrelations were more similar from early to late during a flanker task (*r* = .61) than between rest and the flanker task (*r* = .34) (Kelly, et al., 2008). This is comparable to the values that we observed, and suggests that DN-DAN interactions are dynamically tailored to one's current context. Furthermore, we found that a machine learning classifier was able to distinguish each pair of contexts solely on the basis of DN-DAN connectivity patterns. While the classifier's ability to distinguish cognitive states in the current study was noticeably less accurate than results obtained in other studies using whole-brain connectivity patterns (Gonzalez-Castillo, et al., 2015; Milazzo, et al., 2014; Shirer, et al., 2012), it is quite remarkable that patterns of anticorrelations are sufficiently distinct in each context to allow for above chance-level classification. Together, these findings suggest that DN-DAN connectivity dynamically reconfigures in each cognitive context, thus emphasizing flexibility rather than stability in the relationship between these networks. This implies that DN-DAN interactions during rest do not necessarily reflect the nature of interactions between these networks in general, and cautions against making conclusions about group differences in anticorrelations strictly based on resting state data. Individual and group differences in anticorrelations could potentially reflect differences in mental state rather than fundamental differences in brain function, although parallel age-related reductions in anticorrelation during task and rest have been observed (Spreng, et al., in press).

While anticorrelation may be a frequently occurring state of network organization, particularly for DAN-Core interactions, our findings provide unequivocal support for the idea that current task demands can easily enhance or weaken anticorrelations. In fact, we found that some DN-DAN nodes demonstrated no anticorrelation or even positive connectivity in some cognitive contexts, and during a considerable percentage of windows during rest. The current results suggest that *departures from anticorrelation are a typical phenomenon*, and not necessarily indicative of maladaptive processing. Specifically, it appears that the DN and DAN may alternate between periods of segregation (modular processing) and periods of integration. Consistent with this, prior studies have reported that anticorrelations involving the DN vary across time (E. Allen, et al., 2014; Chang & Glover, 2010) and global brain dynamics also exhibit shifts between periods segregation and integration (Liegeois, et al., 2015; Shine, Koyejo, & Poldrack, 2016). The idea that the DN and DAN may not be not strictly antagonistic is consistent with a large body of work indicating that the relationship between internal and external processing is context dependant (Dixon, et al., 2014). Moreover, given that the variability of mind-wandering is predictive of enhanced externally-oriented task performance (i.e., error awareness), this suggests a complex relationship between internal and external processing (M. Allen et al., 2013). In sum, DN-DAN interactions exhibit substantial temporal and contextual variability, suggesting that their relationship is influenced by changing mental states.

**FPCN and Anticorrelations**. We found that variation in the strength of anticorrelations was not random, but rather, systematically related to FPCN connectivity. Within each context, periods of time characterized by greater negative coupling between the FPCN and DAN, and to some extent greater positive coupling between the FPCN and DN Core, were associated with stronger DAN-Core anticorrelation. Given the opposing nature of these relationships, it seems unlikely that they could be due to noise or a general factor (e.g., arousal). A more likely possibility is that *shifting attentional priorities* encoded by the FPCN exert a top-down influence on anticorrelations.Given that we necessarily used a limited range of tasks, it is possible that different network relationships could emerge in other contexts (e.g., perhaps greater positive FPCN-DAN coupling would be associated with stronger DAN-Core anticorrelations during a visuospatial working memory task). However, the important point is that our findings demonstrate a heretofore unrecognized relationship between FPCN connectivity and the strength of anticorrelations.

While acknowledging that our analyses do not speak to causality, when considered in light of the broader literature on the FPCN, they are consistent with the possibility of a top-down modulatory role over DN-DAN interactions. Abundant evidence suggests that the FPCN encodes task demands, and transmits signals about the current relevance of stimuli, actions, and outcomes to other regions, thus coordinating processing across wide swaths of the cortex (Buschman & Miller, 2007; Cole, Ito, & Braver, 2015; Crowe et al., 2013; Dixon & Christoff, 2012, 2014; Duncan, 2010; Miller & Cohen, 2001; Tomita, Ohbayashi, Nakahara, Hasegawa, & Miyashita, 1999). Consistent with a top-down role, FPCN connectivity with other networks is flexible and adapts in line with task demands (Braun, et al., 2015; Cole, et al., 2013; Gao & Lin, 2012; Spreng, et al., 2010). Here, we extend these findings by demonstrating that FPCN connectivity patterns are tightly coupled with the strength of DN-DAN anticorrelation. One possibility is that the FPCN modulates DN-DAN dynamics based on the extent to which perceptual and conceptual/self-referential processes are currently needed to meet one's goals (Dixon, et al.,2014; Gao & Lin, 2012; Smallwood, et al., 2012; Spreng, et al., 2013; Spreng, et al., 2010; Vincent, et al., 2008). Thus, variability of anticorrelations across time and context may reflect moment-to-moment and context-to-context shifts in the balance of external versus internal information within the focus of attention—and this may be driven by the FPCN.

**Limitations**. A limitation of the current study is that we lack information about the nature and timing of ongoing cognitive activity, and how it relates to the variability of anticorrelations. Future studies could benefit from using online experience sampling to determine the precise cognitive state of each participant as it evolves across time (Christoff, 2012; Fazelpour & Thompson, 2014). Additionally, experimenter controlled variations in task demands on the scale of tens of seconds could also be useful in linking connectivity patterns to mental states (Gonzalez-Castillo, et al., 2015). The present study is also unable to establish whether the FPCN plays a causal role in modulating anticorrelations. This could be addressed in future work by perturbing FPCN functioning via TMS and monitoring the impact on DN-DAN connectivity. Although we have characterized DN-DAN interactions in relation to well-established network boundaries (Yeo, et al., 2011), there is variance in network organization across individuals (Mueller et al., 2013), and hence, future work could improve precision by using individually-tailored network ROIs (Wang et al., 2015). Finally, it could be argued that the contextual variation in anticorrelations that we observed was due to idiosyncratic numbers of attentional lapses in each context. However, several factors make this very unlikely. First, and foremost, the effect of context was not uniform across all DN-DAN connections. For example, from rest to the movie condition, some DN-DAN connections exhibited increased anticorrelation, others exhibited reduced anticorrelation, and some connections exhibited no change. This finding is inconsistent with a general, non-specific factor such as attention/arousal driving the effect of context on anticorrelations. Second, participants reported high levels of attention during the conditions requiring an external focus (range:5.83-6.41 on a 7-point scale, from 1 = not at all paying attention to 7 = paying very much attention). Finally, the machine learning classifier was able to accurately discriminate mental states for each participant based on the data from other participants, implying that there was structure in how anticorrelations varied across contexts. Thus, changes in DN-DAN connectivity across contexts appear to be specifically related to differences in the required cognitive demands.

**Conclusions**. To summarize, DN-DAN interactions are more variable than previously appreciated, suggesting that these networks and the functions they support are not strictly competitive. The DAN exhibits distinct interactions with the three DN subsystems. Furthermore, anticorrelations are not stable, but rather, exhibit a surprising degree of flexibility, increasing and decreasing across time and different cognitive states. Finally, we found a systematic relationship between FPCN-DAN connectivity and anticorrelations, consistent with a possible role for the FPCN in regulating the strength of anticorrelations based on current goals.

## Experimental Procedures

**Participants**. Participants were 24 healthy adults (Mean age = 30.33, SD = 4.80; 10 female; 22 right handed), with no history of head trauma or psychological conditions. This study was approved by the UBC clinical research ethics board, and all participants provided written informed consent, and received payment ($20/hour) for their participation.

**Experimental Conditions**. Each participant performed six conditions in separate six-minute fMRI runs (see **Supplemental Experimental Procedures**):(1) *Resting state*. Participants lay in the scanner with their eyes closed and were instructed to relax and stay awake, and to allow their thoughts to flow naturally. (2) *Movie watching*. Participants watched a clip from the movie “Star Wars:Return of the Jedi”, during which Luke Skywalker engages in a light-saber duel with Darth Vader. (3) *Artwork analysis*. Participants viewed four pieces of pre-selected artwork, each for 90 seconds, and were instructed to attend to the perceptual details and the personal meaning of the art.(4) *Shopping task*. Participants viewed a pre-recorded video shot from a first-person perspective of items within several stores in a shopping mall, and were instructed to imagine that they were shopping for a birthday gift for a friend, and to think about whether each item would be a suitable gift based on their friend's preferences. (5) *Evaluation-based* introspection. Participants reflected on a mildly upsetting issue involving a specific person in their life and were asked to analyze why the situation is upsetting, who caused it, what might happen in the future, and to become fully caught up in their thoughts and emotions. (6) *Acceptance-based* introspection. Participants reflected on a mildly upsetting issue involving a specific person in their life and were asked to cultivate a present-centered awareness, grounded in the acceptance of moment-to-moment viscero-somatic sensations (i.e., to notice and experience arising thoughts, emotions, and bodily sensations with acceptance, and without any elaborative mental analysis or judgment).

**fMRI Data Acquisition**. fMRI data were collected using a 3.0-Tesla Philips Intera MRI scanner (Best, Netherlands) with an 8-channel phased array head coil with parallel imaging capability (SENSE). Head movement was restricted using foam padding around the head. T2*-weighted functional images were acquired parallel to the anterior commissure/posterior commissure (AC/PC) line using a single shot gradient echo-planar sequence (repetition time, TR = 2 s; TE = 30 ms; flip angle, FA = 90°; field of view, FOV = 240 mm; matrix size = 80 × 80; SENSE factor = 1.0). Thirty-six interleaved axial slices covering the whole brain were acquired (3-mm thick with 1-mm skip). Each session was six minutes in length, during which 180 functional volumes were acquired. Data collected during the first 4 TRs were discarded to allow for T1 equilibration effects. Before functional imaging, a high resolution T1-weighted structural image was acquired (170 axial slices; TR = 7.7 ms; TE = 3.6 ms; FOV = 256 mm; matrix size = 256 × 256; voxel size = 1 × 1 × 1 mm; FA = 8°). Total scan time was ∼ 60 minutes. Head motion was minimized using a pillow, and scanner noise was minimized with earplugs.

**Preprocessing**. Full details are provided in **Supplemental Experimental Procedures**. Standard preprocessing steps were conducted with SPM8, including slice-timing correction, realignment (using a 6 parameter rigid body transformation), coregistration with the structural image, normalization to the MNI 152 atlas, and spatial smoothing). Global signal regression was not performed, as it may induce spurious anticorrelations. Additional sources of noise were estimated and regressed out using”CONN“software (Whitfield-Gabrieli & Nieto-Castanon, 2012). Eroded white matter (WM) and CSF masks were used as noise ROIs. Signals from the WM and CSF noise ROIs were extracted from the unsmoothed functional volumes to avoid risk of contaminating WM and CSF signals with gray matter signals. The following nuisance variables were regressed out:three principal components of the signals from the WM and CSF noise ROIs; head motion parameters (three rotation and three translation parameters) along with their first-order temporal derivatives; artifact outlier images; linear trends. A band-pass filter (0.009 Hz <f <0.10 Hz) was simultaneously applied to the BOLD time series during this step.

**Seed-based voxel analyses**. The timeseries of all voxels within the ROIs were averaged, and then first-level correlation maps were produced by computing the Pearson correlation between that seed timeseries and the timeseries of all other voxels. Correlation coefficients were converted to normally distributed z scores using the Fisher transformation to allow for second-level GLM analyses. Results were visualized with CARET brain mapping software (http://brainmap.wustl.edu/caret; Van Essen, 2005; Van Essen et al., 2001).

**Similarity Analysis**. For each participant we extracted DN-DAN node-to-node connections (excluding interhemispheric connections), applied a Fisher r-to-z transformation, and then calculated the correlation between the vectors of connectivity values for each pair of contexts, as well as for the first half (first 3 minutes) and second half (last 3 minutes) of each context. We then computed the average across-context similarity and average within-context similarity on Fisher-transformed correlations. Importantly, we computed similarity for each participant separately (rather than on group averaged connectivity matrices), and then determined average similarity across the group, thus accounting for individual variability.

**Classifier analysis**. To determine whether pairs of conditions could be correctly classified based on patterns of anticorrelations, we used a SVM classifier implemented with The Spider toolbox (Weston, Elisseeff, BakIr, & Sinz, 2005). Following prior work (Dosenbach et al., 2010), we set the cost parameter, C, to 1, and used a radial basis function (RBF) kernel, with sigma set to 2 (similar results were obtained with a linear classifier). For each individual we created a vector consisting of all DN-DAN z-transformed correlations (excluding inter-hemispheric connections) for each condition. The between-network correlations served as input features (96 in total), and were assigned a value of 1 or –1 to specify the condition to which they belonged. We tested the accuracy of the classifier using leave-one-out cross validation:the classifier was trained on the anticorrelation patterns for all but one participant, and then tested on that left-out participant, and this was repeated for each individual. For statistical testing, we obtained an empirical null distribution by performing the classification analysis 1000 times with condition labels randomly permuted. The mean classification accuracy over the 1000 iterations ranged from 49.62% to 50.43% with a standard deviation that ranged from 6.03% to 6.73%, depending on the specific pair of conditions. In each case, inspection of the null distribution revealed that 95% of these models had accuracies below 60.4%. Classification accuracies larger than the 95th percentile of the null distribution were considered to be statistically significant atp <.05.

**Dynamic FC analysis**:To examine time-dependent changes in connectivity, we used a sliding window of 60 seconds, shifted by one timepoint (2 seconds) each time. To limit the possibility of detecting spurious temporal fluctuations in connectivity, we bandpass filtered the data (0.0167 Hz <f <0.10 Hz) such that frequencies lower than 1/w were removed, where w is the width of the window (Leonardi & Van De Ville, 2015).

## Author Contributions

Conceptualization, M.L.D and K.C; Methodology, M.L.D and K.C; Investigation, M.L.D.; Discussion of Analyses, all authors; Writing − Original Draft, M.L.D.; Writing − Review & Editing, all authors. Supervision: K.C.

## Acknowledgments

K.C. is supported by NSERC (RGPIN 327317-11) and CIHR (MOP-115197) grants. J.R.A-H is supported by a”Prospective Psychology Stage 2:A Research Competition“grant from the Templeton Foundation and a Young Investigator Award from the Brain & Behavioral Research Foundation.

